# GlycoSHIELD: a versatile pipeline to assess glycan impact on protein structures

**DOI:** 10.1101/2021.08.04.455134

**Authors:** Michael Gecht, Sören von Bülow, Camille Penet, Gerhard Hummer, Cyril Hanus, Mateusz Sikora

## Abstract

More than 75% of surface and secreted proteins are modified by covalent addition of complex sugars through N- and O-glycosylation. Unlike proteins, glycans do not typically adopt specific secondary structures and remain very mobile, influencing protein dynamics and interactions with other molecules. Glycan conformational freedom impairs complete structural elucidation of glycoproteins. Computer simulations may be used to model glycan structure and dynamics. However, such simulations typically require thousands of computing hours on specialized supercomputers, thus limiting routine use. Here, we describe a reductionist method that can be implemented on personal computers to graft ensembles of realistic glycan conformers onto static protein structures in a matter of minutes. Using this open-source pipeline, we reconstructed the full glycan cover of SARS-CoV-2 Spike protein (S-protein) and a human GABAA receptor. Focusing on S-protein, we show that GlycoSHIELD recapitulates key features of extended simulations of the glycosylated protein, including epitope masking, and provides new mechanistic insights on N-glycan impact on protein structural dynamics.

## Introduction

An estimated 60% of drugs that are currently available or under development targets surface proteins (1). In their cellular context, the overwhelming majority of these proteins are N-glycosylated (2). As shown by atomistic simulations of glycoproteins, glycans are much more dynamic than folded polypeptides (3, 4). Glycans sample extensive arrays of conformations over tens of nanoseconds, hence creating molecular shields that mask large areas of protein surface. While such simulations provide key information on glycan impact on protein structure and interactions with drugs and other biomolecules (5), they require both expert knowledge and extended computing times on specialized supercomputers, limiting their use for routine evaluation of new glycoprotein structures.

Depth and coverage of glycoproteomics analyses are rapidly expanding and a wealth of data is now available on the N- and O-glycomes of a variety of organisms and tissues (6, 7). There is thus a need for reliable and simple tools to evaluate how glycan diversity impacts protein structure and shielding. Here, we generated a library of plausible conformers for a variety of O- and N-glycan types through molecular dynamics simulations (MDS) and developed an open-source pipeline, GlycoSHIELD, enabling non-expert users to graft glycan conformer arrays onto any static protein structures to visualise and analyse the span and impact of diverse glycan shields on protein surfaces.

## Results

### Rationale and validation

As shown by experimental data (8, 9) and simulations (3–5) for SARS-Cov-2 Spike protein (S-protein) and other viral proteins, glycan conformations are sterically constrained by local protein structure and strongly differ from one glycosylation site to another. Accessible glycan conformations are rapidly sampled through thermal agitation, depending mostly on non-specific self-interactions within glycans and extensive interactions with the solvent (10). Excepting the case of specialized glycan binding protein domains (11, 12), glycan interactions with the host protein surface are in most cases non-specific and only transient (4, 13). We thus reasoned that these conformations could be reliably captured by simulations of individual glycans whose conformers may then be grafted to specific glycosylation sites and selected or rejected based on steric hindrance by the protein or cellular membranes. Grafting with steric exclusion has successfully been used to model disordered proteins (14) and reconstruct small N-glycans from individual disaccharides (15). Here we generated a library of 11 N- and O-glycans of high prevalence and particular physiological relevance (2). For each of these glycans we performed extensive (≥3 μs) MDS in solution and obtained large conformation arrays for grafting on any static protein structure (Figure 1A, Supplemental Table S1 and Figure S1, see Methods for details).

**Figure 1.**
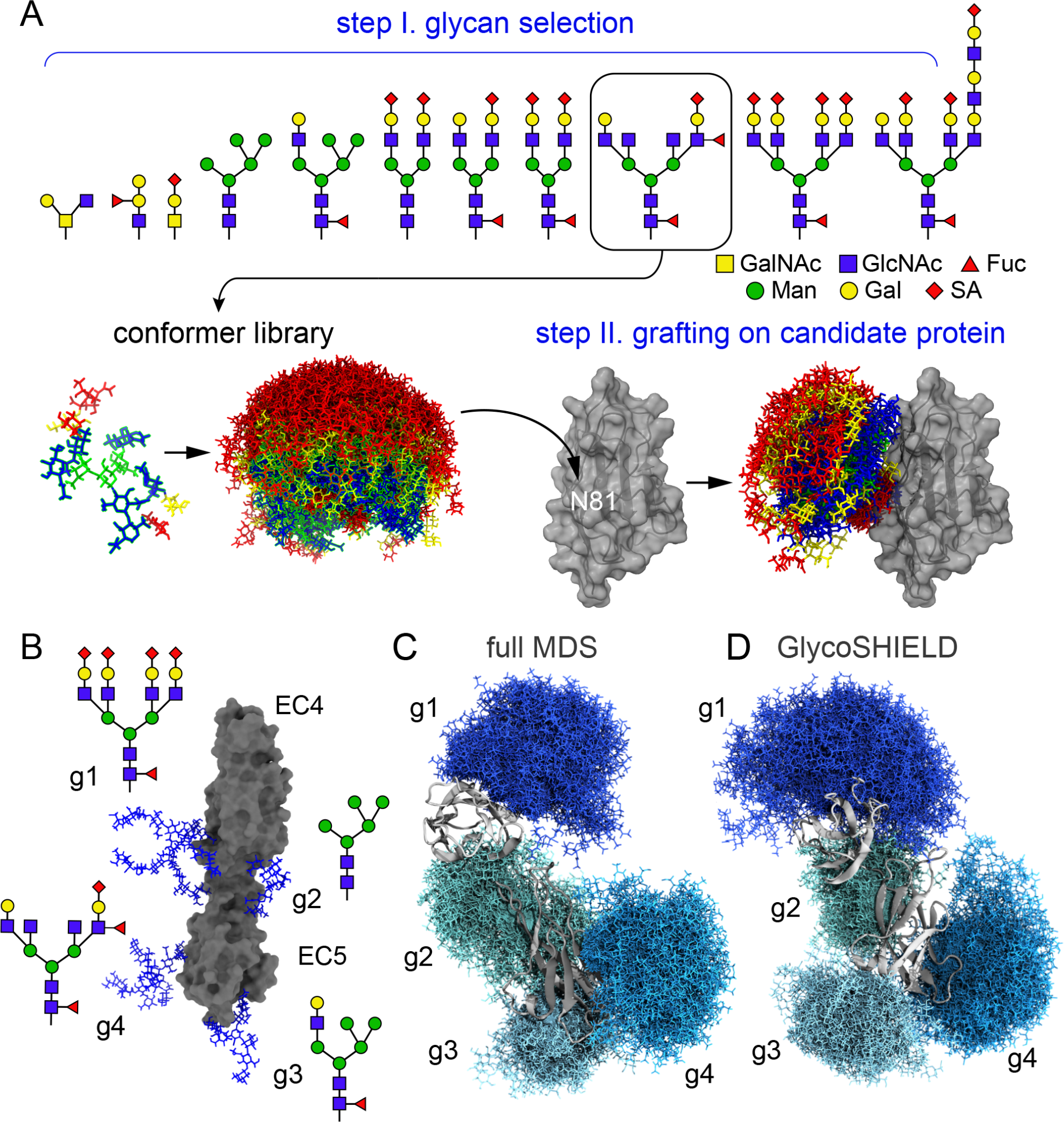
GlycoSHIELD generates realistic glycan shields. **A**) Overview of the pipeline: user provides a 3D protein structure with defined glycosites where glycans from the library of conformers not clashing with the protein are grafted and exported for visualization and analysis (GalNac: N-Acetylgalactosamine, GlcNac: N-Acetylglucosamine, Fuc: fucose, Man: mannose, Gal: galactose, SA: sialic acid). **B**) Structure of N-cadherin EC4-EC5 model system with four distinct N-glycans as indicated at each glycosylation site (g1-g4). **C) -D)** Glycan conformers generated by full MDS (C) or with GlycoSHIELD (D) after alignment on EC4-EC5. Note the comparable morphology and span of the glycan shields obtained by the two approaches.

Glycan dynamics seems to occur primarily without stable interactions with the host protein surface (13, 16). However, labile glycan-protein interactions may still influence the overall morphology of glycan shields and the conformation of the protein. To quantify these effects, we turned to N-cadherin, an adhesion glycoprotein comprising five compact and stable domains (EC1-EC5) connected by linkers, which, in the absence of bound calcium enable extensive interdomain movements (17). To sample both glycan and domain dynamics, we performed 1μs MDS of a reduced system composed of EC4-EC5 domains, which was modelled in its unglycosylated form or with four distinct glycans in the absence of calcium (Figure 1B). Over the course of these simulations, glycans remained mobile and probed extensive arrays of conformations (Supplemental Movie S1), covering large swaths of the protein surface (Figures 1C). Next, we applied GlycoSHIELD to graft corresponding glycans onto a static unglycosylated EC4-EC5 structure. The extent and shape of the glycan shields was well reproduced by GlycoSHIELD (Figure 1C), with only small differences seen in the shielding of the protein surface compared to full MDS of the glycosylated protein (Figure 1C-D and Supplemental Figures S2A). Consistent with our previous observations on other systems (4), analysis of hydrogen-bonds (h-bonds) showed that EC4-EC5 glycans interacted mostly with the solvent (11±3, 24±4, 299±12; means±SD for glycan-protein, glycan-glycan and glycan-water h-bonds, respectively). This indicates that small differences in protein shielding derived from GlycoSHIELD and full MDS were not due to direct glycan interactions with the protein and thus likely resulted from the flexibility of the protein surface and steric hindrance between individual glycans, which, by design, are not accounted for in GlycoSHIELD. We thus conclude that GlycoSHIELD accurately captures the morphology and span of glycan shields.

Next, we focused on the influence of glycans on the dynamics of the protein. We found that the internal motion of single EC domains was largely unaffected by the presence of glycans, with glycosylation sites remaining particularly stable (Supplemental Figure S3). Consistent with previous reports (13, 16), this indicates that glycan dynamics has little impact on the intrinsic stability of compact domains, supporting our approach.

In contrast, glycans affected the domain-scale motion of the whole protein. Unglycosylated EC4-EC5 rapidly folded into a relatively compact and stable conformation. In contrast, the glycosylated protein remained extended and flexible (Supplemental Movie S2). We hypothesized that this more open conformation of glycosylated EC4-EC5 resulted from steric hindrance by glycans preventing close interactions between protein domains (17) (Supplemental Figures S4A-B). To address this, we grafted glycans onto individual snapshots from the MDS of unglycosylated EC4-EC5. Quantification of the radius of gyration of the protein backbone showed that EC4-EC5 compactness was highly correlated with glycan rejection rate during grafting (Supplemental Figures S4C-D), providing a convenient proxy for assessing the entropic cost of steric hindrance on protein conformation (Supplemental Figure S4E). This entropic effect depended on the composition of N-glycans (Supplemental Figure S4F), indicating that the conformation of N-cadherin is modulated by the glycosylation status of the protein. Previous structural studies have shown that calcium-dependent stiffness of the protein is required for strong cadherin-cadherin adhesion (17). Glycans may thus be important to maintain the protein in an elongated conformation in the absence of calcium.

Altogether, these results show that GlycoSHIELD provides reliable estimates of the morphology of glycan shields and their impact on protein conformation.

### GlycoSHIELD reveals partial occlusion of the agonist-binding site of an essential neurotransmitter receptor

Although it is now well established that neuronal proteins, notably neurotransmitter receptors, display atypical N-glycosylation profiles (6, 19), we know surprisingly little about the structural and functional impact of atypical N-glycans on these proteins. To fill this gap, we modelled the glycan shields of a human GABAA receptor (20) (Figure 2A), an inhibitory neurotransmitter receptor that is the target of general anaesthetics and whose unglycosylated structure has been extensively characterized (21). The receptor was modelled with GlnNac2-Man5 (hereafter referred to as Man5, see Supplemental Table S1), one of the two most prevalent glycotypes observed for this receptor in the mammalian brain (6)(Supplemental Figure S5). GlycoSHIELD showed that N-glycans occupy a large volume around the receptor and mask a significant fraction of the surface of the protein (Figures 2B-C). Interestingly, predicted glycan shields partially occluded the agonist binding sites (benzamidine, Figure 2D) as well as the vestibule of the pore (Figure 2E), pointing toward an important role for N-glycosylation in regulating the electrophysiological properties of the receptor (22, 23).

**Figure 2.**
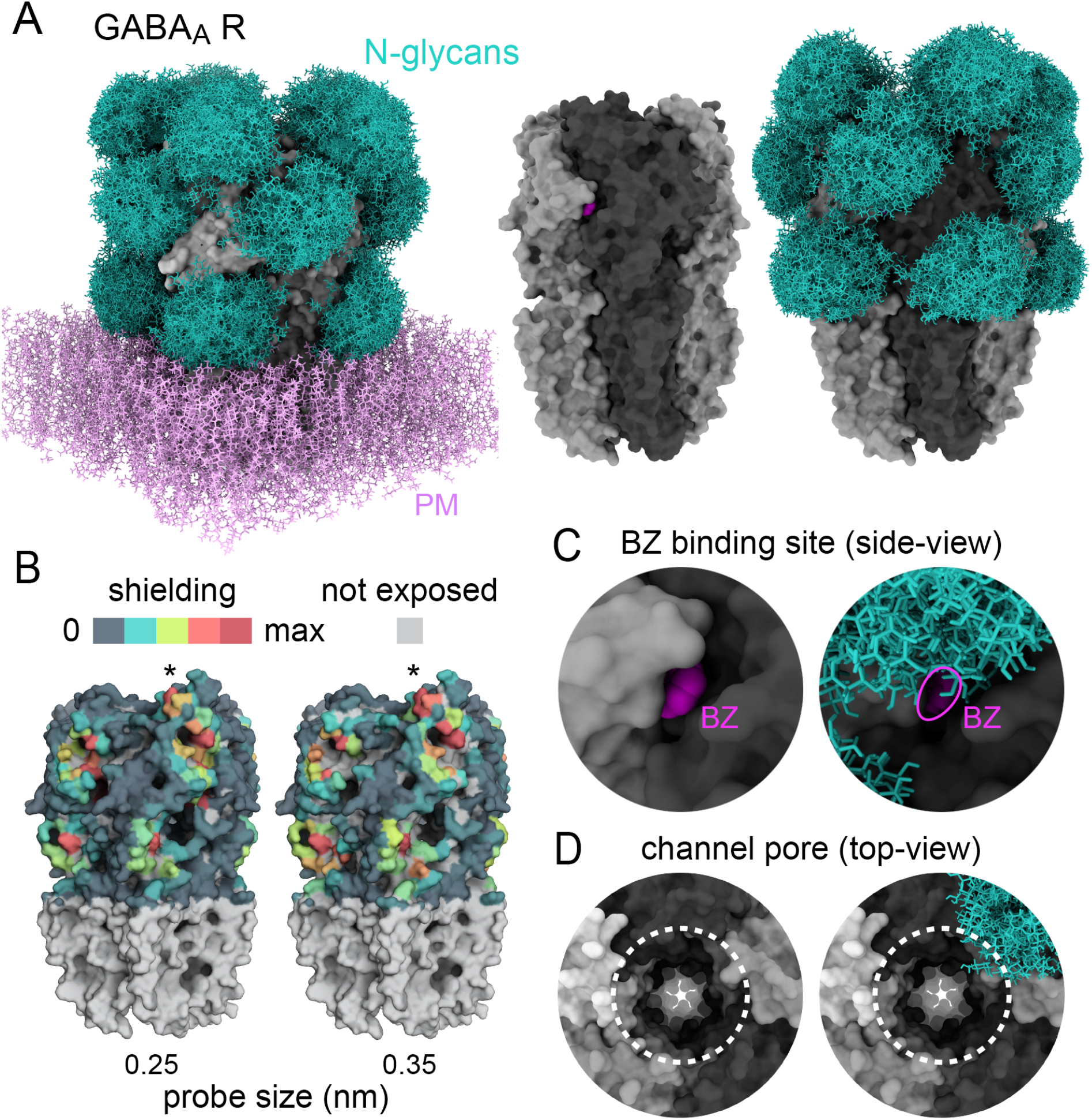
Glycans sculpt the surface of GABA_A_ receptors. **A)** Structure of a human homopentameric GABA_A_ receptor (beta3 subunits, grey) shown with and without patch of plasma membrane (PM, added for visualisation only, pink) with and without reconstituted glycan shields (GlcNac2-Man5, blue in A and B). **B)** 3D-heatmaps of the reduction of solvent accessible surface area (SASA) by glycans for different probe size. Glycan shield on the top of the protein where represented on a single subunit only (asterisk) to better document the impact of single glycans. **C-D)** Magnified views of the agonist binding site with bound ligand (benzamidine or BZ), shown in fuchsia (C) and the extracellular vestibule of the channel pore (dotted lines in D). In C and D, note the partial occlusion of the ligand binding site and the ending of the glycan shield at the opening of the channel pore, respectively. For better visualization of the pore, only a single glycan was represented in C.

Thus, GlycoSHIELD may unravel novel and unrecognised features of glycoprotein structures, providing incentive and useful focal points for extended computational and experimental studies.

### GlycoSHIELD provides new insights on glycan impact on SARS-CoV-2 S-protein dynamics and recognition by the immune system

Extensive glycosylation of type I viral fusion proteins contributes to the success of enveloped viruses in evading the immune system and hinders vaccine and drug development. In this context, the glycan shield of the SARS-CoV-2 S-protein has been a focal point in the current anti-pandemic effort to develop vaccines that can elicit production of neutralizing antibodies directed against existing and emerging S-protein variants (24, 25).

S-protein is a homo-trimer that comprises sixty-six N-glycosylation sites and is heavily glycosylated (9, 26). Glycosylation may strongly vary depending on host cell types and organisms, likely impacting how S-protein interacts with its cellular receptor ACE2 (5) (Angiotensin Converting Enzyme 2) and activates membrane fusion. Large-scale MDS has provided compelling information on the structural impact and epitope-masking properties of specific N-glycans identified on S-protein after expression in common expression systems (4, 9). Due to prohibitive costs in computing resources, this approach is not readily amendable to screening of large arrays of tissue-specific N-glycans, calling for more versatile and less costly options. As a first step in that direction, we used GlycoSHIELD to compare shields generated by various glycans on S-protein. As done for EC4-EC5, we compared glycans shields generated with GlycoSHIELD on the naked protein with those generated by extended simulation of the glycosylated protein. Although the glycans used in GlycoSHIELD and full MDS were not strictly equivalent (Supplemental Figure S6), we found that the morphology of reconstituted shields was in good agreement with shields derived from MDS (Figures 3A)(4), and, as expected, differed depending on glycan composition (Supplemental Figure S7).

**Figure 3.**
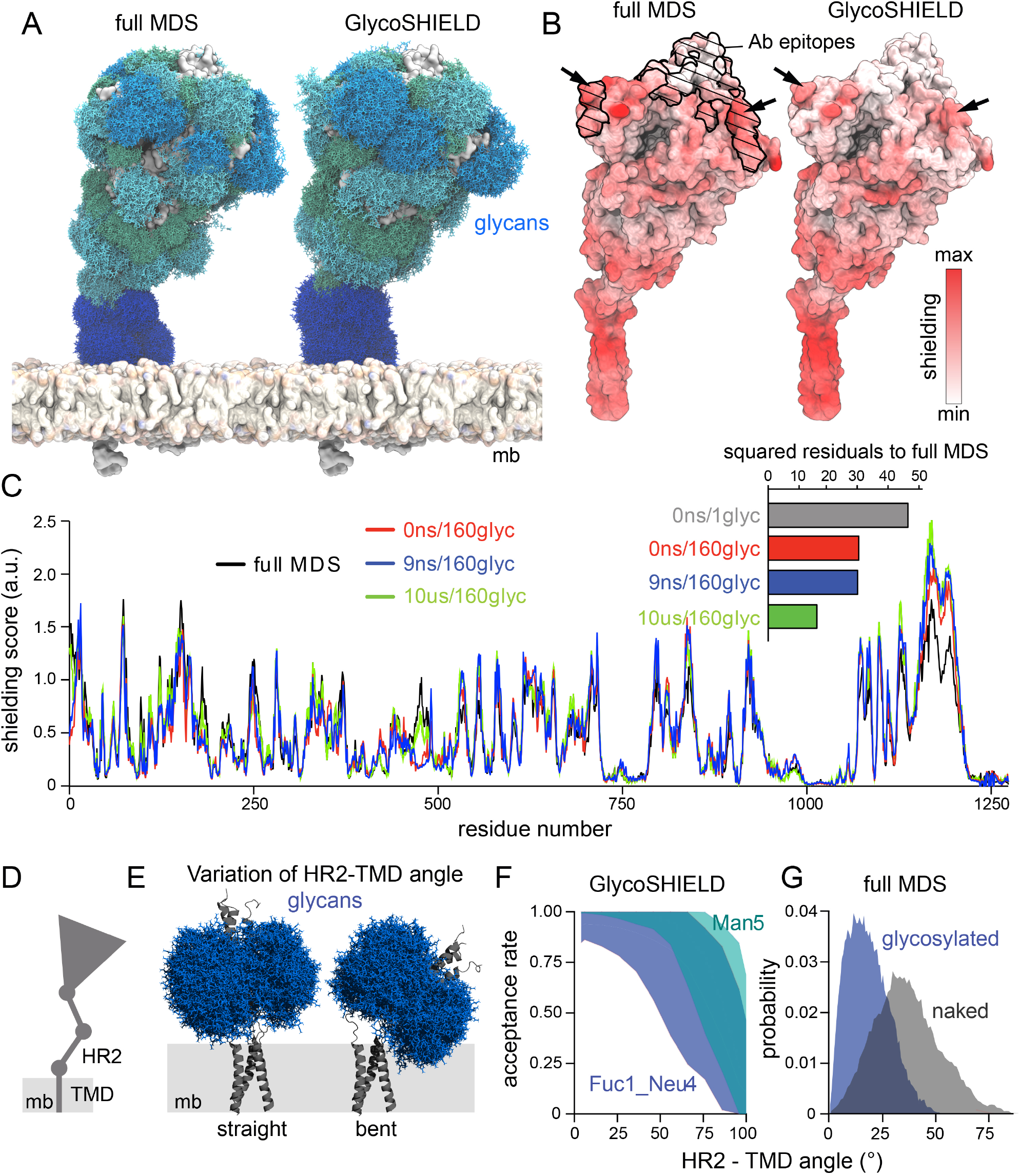
N-glycans may impact the conformation of SARS-Cov-2 Spike protein and its recognition by antibodies. **A**) S-protein glycan conformers sampled from 10 μs of MDS or reconstructed with GlycoSHIELD. In both structures, 160 glycan conformers are displayed for each glycosite. Membranes were added for visualisation purposes. **B)** Glycan shielding of S-protein extracellular domain calculated from MDS versus GlycoSHIELD. Shown are 3D heatmaps of accessibility scores calculated by ray analysis. Higher color intensities indicate higher shielding. Arrows indicate predicted shielded areas within specific antibody epitopes (black lines and hatched areas). **C)** Shielding of S-protein residues calculated from the original 10 μs MDS (black) and recalculated with GlycoSHIELD: for a single protein conformer and single glycan conformer (gray), single protein conformer and 160 glycan conformers (red), 9 protein conformers corresponding to the initial 9 ns of the simulation and 160 glycan conformers (blue) and 100 protein conformers sampled at 100 ns intervals with 160 glycan each. Inset shows corresponding sum of squared residuals of shielding in respect to full MDS. **D)** Scheme of S-protein main domains. **E)** Model of S-protein HR2 and TMD domains in upright and inclined position with glycan shields shown for N1194 (Fuc1_Neu4 glycans, blue). Note the higher proportion of glycan conformers clashing with the membrane (mb, grey band) for the bent protein. **F)** Rate of glycan conformer rejection due to clashes with membrane as a function of HR2 bending with all Fuc1_Neu4 (blue) or all Man5 N-glycans (green). Note the increase of steric constraints upon bending and the distinct impact of the two N-glycan types. **G)** Angle distribution of unglycosylated and glycosylated S-protein stalk in 3.5 μs MDSs showing decreased bending of the glycosylated protein.

As shown by us and other groups, glycan shields may impact the recognition of the S-protein by antibodies (27), thereby affecting host immune responses and the outcome of clinical interventions with neutralizing antibodies. We thus sought to determine whether GlycoSHIELD could capture the masking of S-protein epitope as seen in extended simulations of the glycosylated protein (4). To this end we performed a ray accessibility analysis, i.e., calculated the shading of protein surfaces by glycans upon illumination with randomly oriented light-rays (see Methods). We hence compared different levels of GlycoSHIELD reconstructions on naked (*i.e.* deglycosylated) S-protein conformers extracted from full MDS of the glycosylated protein. As shown by 3D shielding heatmaps, GlycoSHIELD captured almost fully epitope shielding predicted by full MDS (Figure 3B).

We then determined the relative contributions of protein and glycan dynamics to glycan shielding. To this end, we extracted snapshots of S-protein conformation over short and extended segments of the full MDS, specifically 1, 10 and 100 conformers, which were extracted from the first simulation frame (“0 ns”), from the first 9 ns (“9 ns”) of the simulation, or from the entire 10 μs-long simulation (“10 μs”).

We then performed a ray-analysis after grafting 160 glycan conformers (*i.e.*, the smallest number of accepted conformations across all the glycosylation sites of the protein) onto these 1-100 protein conformers (Figure 3C or D). Consistent with 3D shielding heat maps (Figure 3B), shielding scores calculated with GlycoSHIELD on a single S-protein conformer were in good agreement with full MDS and were only slightly improved by sampling multiple protein conformers (Figure 3C). Differences in shielding scores along the protein were apparent in the first 30 N-terminal residues of the protein and in the loops of the receptor-binding domain (RBD, residues 450-500)(Figure 3C). These differences may be attributed to the significant conformational changes that occur in these regions over μs time scales (Supplemental Figure S8) and which are thus only captured by extended simulations. As seen by comparing squared residuals of GlycoSHIELD versus full MDS shielding plots (Figure 3D), the grafting of single glycan conformers onto a static S-protein structure was already sufficient to predict glycan shielding at a coarse level (0 ns/1glyc in Figure 3D). As expected however, grafting arrays of glycan conformers significantly improved shielding prediction (0 ns/160glyc). Averaging over protein conformers sampled over extended durations further improved shielding predictions (10 μs/160glyc). Interestingly however, sampling over short segments of the protein trajectory did not improve the prediction made on a single protein conformer (compare (9 ns/160glyc to 0 ns/160glyc). Thus, these results indicate that GlycoSHIELD generates realistic glycan shields on stable protein structures regardless of short-scale protein dynamics, enabling reliable predictions of the protein surfaces available for interactions with the immune system.

Cryo-electron tomography *in situ* and MDS approaches have shown that S-protein stalk remains very flexible (25), likely affecting S-protein interaction with ACE2 and thus increasing SARS-CoV-2 avidity for target cells (Figure 3E). It is currently not known whether large glycans present on the protein stalk modulate its flexibility. As shown in Figure 3A, large glycans located close to S-protein transmembrane segments may interact with the virus membrane, hence constraining the orientation of the protein. We used GlycoSHIELD to test this hypothesis. To this end, we generated a truncated model comprising S-protein heptad repeat 2 (HR2) region and transmembrane domain (TMD, Figure 3E). To mimic the flexibility of the HR2-TMD hinge, we generated a range of protein conformers, varying the angle between HR2 and TMD and used GlycoSHIELD to quantify glycan clashes with the virus membrane (Figure 3F). We found a steep increase of conformer rejection rate for protein conformers oriented towards the membrane (Figures 3G). This suggests that HR2 glycans favour upright positions of S-protein, consistent with cryoEM data on the native protein (26). To validate this prediction, we simulated the glycosylated and unglycosylated models of truncated S-protein embedded in a realistic membrane (Figure 3G). We found that the glycosylated protein remained more upright than its non-glycosylated counterpart (Figure 3G), thus directly validating predictions made with GlycoSHIELD. Thus, GlycoSHIELD may predict glycan impact on protein structural dynamics.

Altogether, these results show that GlycoSHIELD reliably captures key features of extended MDS of complex systems, enabling predictions on the structural impact of N-glycans on protein surface accessibility and preferred conformation.

## Conclusion

The present study shows that GlycoSHIELD provides a straightforward and accessible approach for obtaining novel and still missing quantitative information on glycoprotein morphology and structural dynamics. This reductionist approach greatly reduces required computing resources, time and technical know-how. Yet, by design, it ignores direct interactions of glycans with the protein and reciprocal interactions between glycans. Glycans modelled with GlycoSHIELD are thus not expected to recapitulate specific glycan-protein binding, as occurring for example for glycan docking on lectins (11, 12) or S-protein (5). Reconstituted shields provide, however, a probabilistic view of glycan positions integrated over the nearly complete ensemble of possible conformations, and as shown here, are very similar to shields predicted by MDS. While GlycoSHIELD cannot substitute for extended MDS of complete systems, it thus fills an important gap by enabling non-expert users to generate reliable models of glycan shields and analyse their masking properties for the overwhelming majority of glycoproteins for which such shields have not been modelled yet.

Tools exist to add single glycan conformers onto existing protein structures (28–30). Yet, these do not fully account for possible glycan conformational space and, when used for system preparation for MDS, may be biased and excessively constrained by initial glycan conformation. Selecting glycan conformers generated by GlycoSHIELD may alleviate this problem, allowing multiple locally equilibrated structures to be used as replicates of a given glycosylated site.

As shown in particular for SARS-CoV-2 and other coronaviruses, naturally occurring mutations may influence virus cellular tropism (31). This may in turn affect virus glycosylation status and recognition by the immune system. The issue is also critical for cancer immunotherapy as cancers are typically associated with aberrant protein glycosylation (32, 33). Versatile tools enabling rapid assessment of the impact of diverse glycans on protein shielding will thus likely be of value in this context. As illustrated by RosettaAntibody Design (34), dynamic structural modelling greatly accelerates the design and generation of custom antibodies. Because of its modular and open-source structure, GlycoSHIELD may easily be integrated into such pipelines, thus adding an important but yet still missing dimension to *in silico* screening of antigens and antibodies.

## Acknowledgments

We thank IDRIS for the allocation of HPC resources on the Jean Zay supercomputer (allocations #2020-AP010711998 and #2021-A0100712343 to CH). We thank the Max Planck Computing and Data Facility for providing computational resources. We thank the French Institute for Bioinformatics (IFB) for their support. MS is supported by a Fonds zur Förderung der wissenschaftlichen Forschung Schrödinger fellowship (J4332-B28). Work in the laboratory of CH is supported by the Agence Nationale de la Recherche (grants #ANR-16-CE16-0009-01 and #ANR-21-CE16-xxxx-xx) and the Foundation NRJ.

## Author contributions

MG, SVB and CP performed analyses and edited the manuscript.

GH and CH provided resources and data and edited the manuscript.

CH and MS designed the project, performed analyses and wrote the manuscript.

## Conflict of interest statement

The authors have no conflict of interest to declare.

## ONLINE METHODS

### Molecular dynamics simulations (MDS)

#### System preparation

Models used for MDS were solvated, glycosylated and inserted in membrane models with CHARMM-GUI (1), unless mentioned otherwise. Systems were solvated using TIP3P water model (2) in the presence of 150mM NaCl and were prepared for simulations with CHARMM36m protein (3) and glycan (4) force fields.

##### Single glycan systems

Individual N- and O-glycans (Supplemental Table S1) were attached to the central residue of neutralised GLY-ASN-GLY (N-glycans) or ALA-THR-ALA tripeptides (O-glycans) and placed in enlarged (by 15-20 Å) rectangular simulation boxes to prevent self-interactions.

##### EC4-EC5

EC4-EC5 domains (residues 331-542) were derived from N-cadherin structure PDB-ID 3q2w (5) and were prepared with neutral N and C-termini, in a unglycosylated and a glycosylated form with the N-glycans shown in Figure 1B.

##### Full length glycosylated SARS-CoV-2 Spike protein (S-protein)

The preparation of this system has been described elsewhere (6) and experimentally validated by cryo-electron tomography (7). In brief, the system contained four membrane-embedded S-protein trimers assembled from available structures and *de novo* models for the missing parts. S-protein head was modelled based on PDB ID: 6VSB (8) with one RBD domain in an open conformation. The stalk connecting S-protein head to the membrane was modelled *de novo* as two trimeric coiled coils (9). The TMD as well as the cytosolic domain were modelled *de novo*. Glycans were modelled according to glycoproteomics data (10) (Supplemental Figure S6 for glycan composition).

##### S-protein stalk (HR2-TMD)

A truncated system comprising the heptad repeat region 2 (HR2) and the palmitoylated transmembrane domains (TMD) of S-protein (residues 1161-1273) was generated from the structure of the unglycosylated or the glycosylated forms of the full length S-protein model described above. N-termini were neutralized and the protein was inserted in a hexagonal patch of plasma membrane (lipid ratios for inner leaflet: POPC:POPE:POPS:CHOL 35:2:20:20; lipid ratios for outer leaflet: POPC:DPSM:CHOL 1:1:1), with the membrane oriented in the xy plane and TMD oriented along the Z-axis of the simulation box.

#### Minimization, equilibration and production

MDS were performed with GROMACS 2018.2, 2019.6 or 2020.2 engines (11). Potential energy of each system was first minimized (steepest descent algorithm, 5000 steps) and subjected to equilibration procedures (see below). Cutoff radii for non-bonded interactions were applied at 1.2 nm, with force switch at 1 nm for van der Waals interactions. Particle Mesh Ewald algorithm was used for long-range electrostatic interactions. All simulations were performed at ambient pressure and 310 K temperature. Atom positions were stored at 100 ps intervals.

##### EC4-EC5

The glycosylated and unglycosylated systems were equilibrated in NVT ensemble for 18.75 ps (with 1 fs time-steps using Nose-Hoover thermostat (12)). Atom positions and dihedral angles were restrained during the equilibration, with initial force constants of 400, 40 and 4 kJ/mol/nm^2^ for restraints on backbone positions, side chain positions and dihedral angles, respectively. The force constants were gradually reduced to 0. Systems were additionally equilibrated in NPT ensemble (Parinello-Rahman pressure coupling (13) with the time constant of 5 ps and compressibility of 4.5e^−5^ bar) over the course of 10 ns with a time step of 2 fs. Hydrogen bonds were restrained using LINCS algorithm (14). During the production runs, velocity-rescale thermostat was used (15) and temperature was kept at 310K. MDS of unglycosylated and glycosylated EC4-EC5 were performed for a total duration of 1μs each.

##### S-protein stalk (HR2-TMD)

Preparation and production runs were performed similar to EC4-EC5. Total NVT equilibration was extended to 125 ps, NPT equilibration comprised additional 1 ns using Berendsen barostat (16) (with semi-isotropic coupling, time constant of 5 ps and compressibilities equal to 4.5e^−5^ bar^−1^). The restraint force constants were gradually reduced from 4000, 2000, 1000 to 0 kJ/mol/nm^2^ for backbone positions, side chain positions and dihedral angles, respectively. Unglycosylated and glycosylated HR2-TM systems were each simulated for 3.5μs.

##### Single glycan systems

After minimization, 12.5 ps NVT equilibration was performed using Nose-Hoover thermostat, with all hydrogen bonds restrained. Atom positions and dihedral angles were restrained during the equilibration, with initial force constants of 400, 40 and 4 kJ/mol/nm^2^ for restraints on backbone positions, side chain positions and dihedral angles, respectively. The force constants were gradually reduced to 0. NPT production runs were performed using Parinello-Rahman barostat and velocity-rescale thermostat as described above. Total simulation time per glycan system was between 3 to 4 μs.

### Trajectory analyses

Root-mean-square deviations (RMSD) and root-mean-square-fluctuations (RMSF) of atom positions, gyration radius of the protein backbone, and hydrogen bond numbers were calculated with GROMACS 2020.2 or 2019.6 (11). Angle and distance measurements were performed with MDAnalysis (17). HR2 angle with the membrane normal was calculated by measuring the deflection of a vector defined by the centres of mass of the residues flanking the coiled-coil region (residues 1160 and 1181) from the Z-axis. Principal component analyses were performed with MODE-TASK (18).

### Quantification of protein surface accessibility with ray-analysis

The accessibility of S-protein surfaces was probed by illuminating the protein by diffuse light as described elsewhere (6). In brief, 10^6^ light rays of random orientation were generated from the inner surface of a 25 nm-radius spherical dome centred on the centre of mass of the protein. Rays were considered absorbed when located ≤ 2 Å from an heavy atom. The shielding effect of glycans was quantified by calculating the irradiation of the protein surface with and without including glycans. Representative antibody epitopes were taken from ref. [6].

### Quantification of protein solvent accessible surface area (SASA)

To quantify the impact of glycans on protein surface accessibility, per-residue SASA was calculated in parallel for unglycosylated and glycosylated proteins (SASA_nogly_ and SASA_gly_, respectively) where SASA_gly_ is the accessible surface area of the protein in the presence of glycans. Calculations were performed with GROMACS *gmx sasa* and custom Python scripts (see code description and availability below). In case of multiple glycan conformers, SASA_gly_ was calculated and averaged over all conformations. Shielding was calculated as a relative SASA:

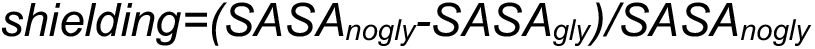

A shielding value of 1 denotes the full shielding of a residue, and 0 denotes no shielding. For glycans grafted on EC4-EC5, SASA was calculated separately for each glycosylation site: trajectories containing the static protein structure and a mobile grafted glycans were used to calculate the average shielding stemming from a specific glycan. The final shielding score of a residue was then defined as the maximum of the shielding scores obtained for each individual glycosylation site. Grafting was performed with 100 ps time step, CG mode and 3.5 Å cutoff, yielding at least 7750 conformers per site, see GlycoSHIELD details below.

To ensure a fair comparison with the grafted system, glycans from EC4-EC5 MDS were locally aligned onto the initial protein conformer, yielding a trajectory containing a static protein and mobile glycans. Shielding was calculated for a random subset of 7750 frames per site.

### GlycoSHIELD

#### Methodology and user scripts

A library of conformers of 11 distinct glycans (up to 40 000 conformers per glycan) was generated and is publicly available on Zenodo (https://zenodo.org/record/5083355). Each conformer contains structural information about the glycan and a tripeptide to which glycan was attached. For convenience, trajectories sampled at 100 ps and 1 ns are provided (GlycoSHIELD time step). The library has an open character and additional glycans species as well as more extensive sets of conformers will be made available in the future. Users are also encouraged to share glycan+tripeptide trajectories from existing glycoprotein simulations.

For every glycosylation site, users select a glycan to be grafted from the available library. GlycoSHIELD aligns every glycan conformer using the backbone atoms of its tripeptide anchor onto the corresponding residues on the protein, i.e., the selected ASN or THR/SER residue ± 1 flanking residue. The selected array (excluding the flanking residues) is then tested for clashes with protein atoms within a threshold radius, and clashing conformers are excluded from the glycan array. Rejection criterion can be based on all protein and glycan atoms or on protein α-carbons and glycan ring-oxygens. The latter, dubbed coarse-grained (CG) threshold, is especially recommended for larger protein systems, where clash search may require extended calculations.

Based on comparison with atomistic simulations (Supplemental Figure S2B), threshold values of 0.7 Å and 3.5 Å are recommended for atomistic and CG clash tests, respectively. In addition, the presence of a biological membrane can be mimicked after orienting the protein with the z-axis perpendicular to the membrane by setting up glycan rejection above or below defined values along the z-axis using z_max_ or below z_min_ thresholds, respectively.

GlycoSHIELD output consists of glycan frames successfully grafted onto the static protein for each probed glycosylation site which are exported in .XTC and .PDB formats. In case of multiple glycosylation sites, the script “GlycoTRAJ.py” may be used to merge this information into a single master trajectory with all grafted glycans. Typically, the smallest number of glycan frames grafted across all glycosylation sites of the protein can be selected, thus generating a “trajectory” with static protein and mobile glycans.

### Code description and availability

Source code and tutorial are available on GitHub (https://github.com/GlycoSHIELD-MD/GlycoSHIELD-0.1). Code for GlycoSHIELD installation *via* conda (e.g., « conda install GlycoSHIELD ») is in preparation.

#### GlycoSHIELD

As an input, user provides a static structure file (PDB format) or series of structures (PDB or GROMACS XTC format). Glycosylation sites and glycans to be grafted are provided by users as batch input files as follows: the first column defines protein chain symbol as defined in PDB, followed by comma-separated glycosylation site residue number together with its two flanking residues, third column defines the tripeptide residue numbers as defined in the glycan conformer library (typically 1,2,3). Fourth and fifth column provide path to the glycan conformer reference structure (PDB) and trajectory files (XTC), respectively. Finally, the last two columns define names of the PDB and XTC output files generated by the script, containing static protein structure and a number of the accepted glycan conformers (see Tutorial page 22).

The syntax of the main GlycoSHIELD is as follows, with optional parameters in square brackets:

~~~
--mode - mode of trimming of the glycan conformer library. CG (coarsegrained): distances between alpha carbons of amino acids and ring oxygens of glycans are calculated to remove clashe, less precise but faster for larger systems. AA: distances between all protein and sugar atoms are calculated to remove clashes.
--threshold - a minimum distance between protein and glycan atoms below which glycan conformer is rejected
--protpdb - structure file containing protein coordinates in PDB format
--protxtc - optional, sequence of conformations of a protein (i.e., a trajectory). Each of the frames will be grafted separately.
--dryrun - does not produce grafted structures, but informs about the number of accepted glycan conformers per site and per protein conformer. Useful, e.g., when entropic penalty for a number of distinct protein conformations is calculated.
--ignorewarn - does not stop the grafting if no glycan conformers not clashing with the protein were found.
--zmax and --zmin - force rejection of glycans that extend beyond these values (in angstrom) along the Z axis. This option is useful for mimicking steric hindrance by cellular membranes.
--shuffle-sugar - by definition glycan conformers are taken from the library in a sequential fashion. This option introduces additional randomization by shuffling the order of the library of conformers for every glycosite.
--help - displays a help message
~~~

#### GlycoSASA

Shielding calculation requires a set of glycan trajectories generated by the GlycoSHIELD script (see above), each containing the same, static protein conformer, and a mobile glycan. Script syntax is as follows:

~~~
--pdblist - comma-separated list of paths to the PDB reference files
--xtclist - comma-separated list of paths to the XTC trajectories
--probelist - comma-separated list of the probe radii, for which SASA should be calculated (in mm)
--plottrace - boolean, should a plot of per-residue SASA be also produced
--keepoutput - whether the temporary output files of the SASA calculation should be kept. For debugging purposes
--ndots - how many dots should be used per atom in the SASA algorithm (see GROMACS manual for the details)
--mode - by default shielding is calculated as a maximum (max) of the shieldings at each glycosylation site. User can choose to calculate also the average shielding (avg)
--endframe - the last frame to read for glycan trajectories, assumes 1ps time step
~~~

Output consists of a TXT file that assigns a shielding value to each residue. These values (multiplied by 100) are also written into the b-factor column of the PDB file. Residues that are not visible to the probe regardless of glycosylation indicated are further marked by a value of 0 in the occupancy column of the PDB file. The output files are named: maxResidueSASA_probe_RPROBE.txt and maxResidueSASA_probe_RPROBE.pdb, where RPROBE is a probe radius. If per-residue shielding plot was requested (--plottrace), a plot is created “under maxResidueSASA_probe_RPROBE.pdf”. Currently, only single chain PDBs are supported by GlycoSASA.py

### Dependencies

GlycoSHIELD requires the MDAnalysis-1.1.1 (17), numpy-1.21.1 (19) and Matplotlib-2.2.3 (20) Python packages. Calculation of the glycan-dependent Solvent Accessible Surface Area (SASA) requires GROMACS-2019.2 (gmx sasa tool, only tested by us in version 2019.2).

### Glycan shield modelling with GlycoSHIELD

#### EC4-EC5

Unglycosylated and glycosylated EC4-EC4 trajectories were aligned to select similar protein conformers from the two simulations. For comparison of shields derived from actual MDS and GlycoSHIELD, all the glycan conformers from the simulation of the glycosylated protein were locally aligned to the selected protein conformer. The unglycosylated protein was glycosylated with GlycoSHIELD with the same glycans than the glycosylated protein, with a CG distance threshold of 3.5 Å.

#### GABA_A_ receptor

The protein model was generated from the structure of a human homopentameric GABA_A_ receptor, PDB-ID: 4COF (21). The 11 accessible N-glycosylation sites of the protein structure were then glycosylated with Man5 and glycan shields modelled with an “AA” threshold of 0.8 Å and taking into account steric hindrance by the upper leaflet of the membrane using GlycoSHIELD --zmax option.

#### Full-length SARS-CoV-2 Spike protein

Grafting was performed on a full-length S-protein trimer, either using the heterogeneous glycans as shown in Supplemental Figure S6 or a homogeneous coverage with Man5 or Fuc1_Neu4 (see glycan definitions in Supplemental Table S1).

For the ray analysis, heterogeneous grafting was further performed on the first 10 ns of the full S-protein simulation at 1 ns intervals, resulting in 160 glycan conformers grafted on each glycosylation site and each protein conformation (CG mode with 3.5 Å threshold, glycans grafted at 100 ps intervals).

#### S-protein stalk (HR2-TMD)

Models of the S-protein stalk with different bending angles were generated from unglycosylated HR2-TMD as follows. The linker region from the HR2-TMD system (residues 1205-1211) was removed. TMD was oriented along the Z-axis and HR2 placed above it at the correct distance. Next, HR2 was rotated by varying the angle between the Z-axis and a vector defined by the centres of mass of HR2 N-terminal residues, with the point of rotation fixed above TMD N-terminal residues. Structures corresponding to a range of angles were generated and linker region reconstructed using MODELLER (22). Each of the structures was then processed with GlycoSHIELD to reconstruct glycan shields with Man5 or Fuc1_Neu4 for all glycosylation sites. (CG mode, 3.5 A threshold, glycans grafted at 1ns intervals).

### 3D structure rendering

Rendering of molecular structures were performed with PyMOL (23), VMD (24) or ChimeraX (25). Glycan conformers were subsampled for rendering of representative conformations on displayed renders.

## Supplementary Figures

**Supplemental Table S1.**
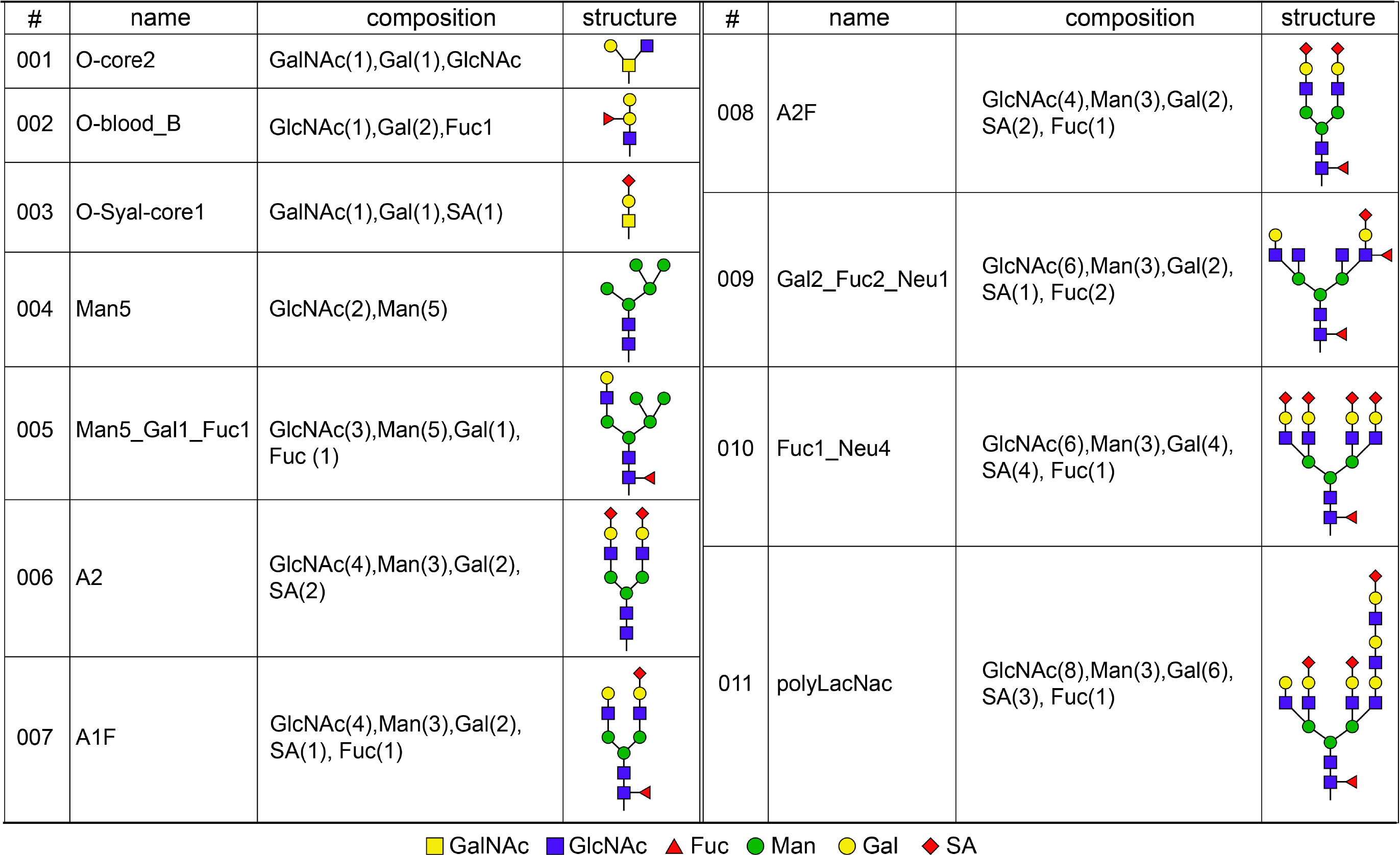
GlycoSHIELD glycan library 1.1 Name, structure and type of glycans currently available in GlycoSHIELD library

| # | name | composition | structure | # | name | composition | structure |
| --- | --- | --- | --- | --- | --- | --- | --- |
| 001 | O-core2 | GalNAc(1),Gal(1),GlcNAc |  | 008 | A2F | GlcNAc(4),Man(3),Gal(2),SA(2), Fuc(1) |  |
| 002 | O-blood_B | GlcNAc(1),Gal(2),Fuc1 |  |  |  |  |  |
| 003 | O-Syal-core1 | GalNAc(1),Gal(1),SA(1) |  |  |  |  |  |
| 004 | Man5 | GlcNAc(2),Man(5) |  | 009 | Gal2_Fuc2_Neu1 | GlcNAc(6),Man(3),Gal(2),SA(1), Fuc(2) |  |
| 005 | Man5_Gal1_Fuc1 | GlcNAc(3),Man(5),Gal(1),Fuc (1) |  | 010 | Fuc1_Neu4 | GlcNAc(6),Man(3),Gal(4),SA(4), Fuc(1) |  |
| 006 | A2 | GlcNAc(4),Man(3),Gal(2),SA(2) |  | 011 | polyLacNac | GlcNAc(8),Man(3),Gal(6),SA(3), Fuc(1) |  |
| 007 | A1F | GlcNAc(4),Man(3),Gal(2),SA(1), Fuc(1) |  |  |  |  |  |

■ GalNAc ■ GlcNAc ▲ Fuc ● Man ● Gal ◆ SA

**Supplemental Figure S1.**
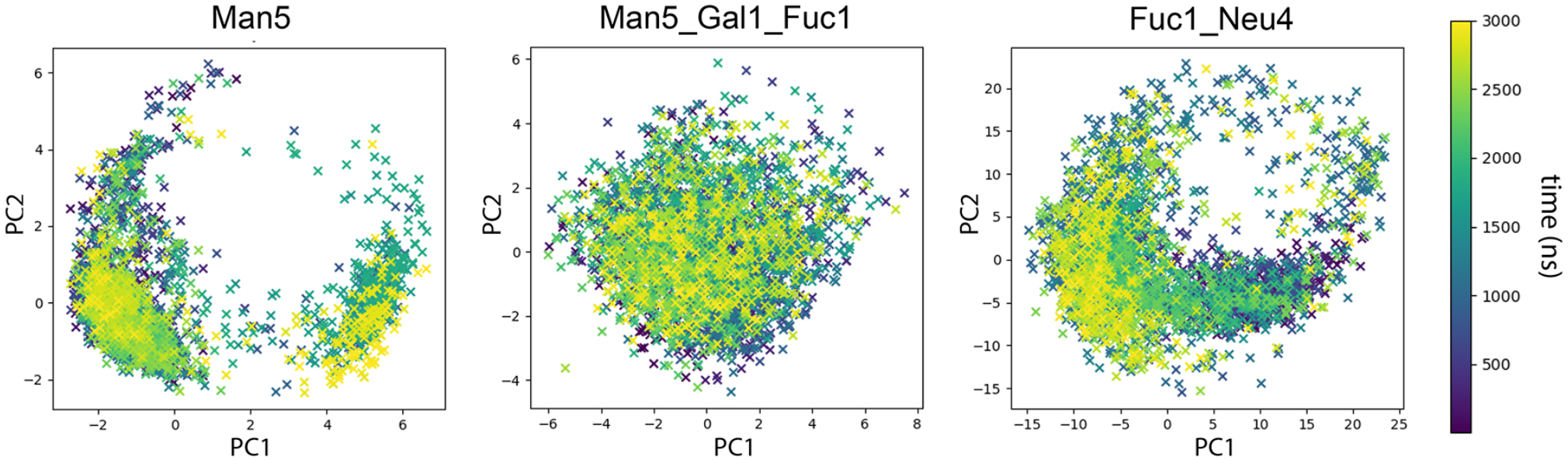
Characterization of conformation sampling by principal component analysis. Variation of PCA2 = f (PCA1) over time showing evolution of glycan atom positions over simulated trajectories for generic examples of high-mannose (Man5), hybrid (Man5_Gal1_Fuc1) and complex (Fuc1_Neu4) N-glycans. Note that conformation clusters are revisited multiple times over the course of the simulations.

**Supplemental Movie S1. MDS trajectories of unglycosylated and glycosylated EC4-EC5 dynamics.** Protein backbones, and glycans are shown in grey and blue, respectively. Time in ns. Unglycosylated and glycosylated EC4-EC5 are shown on the left and the right panel, respectively.

**Supplemental Figure S2.**
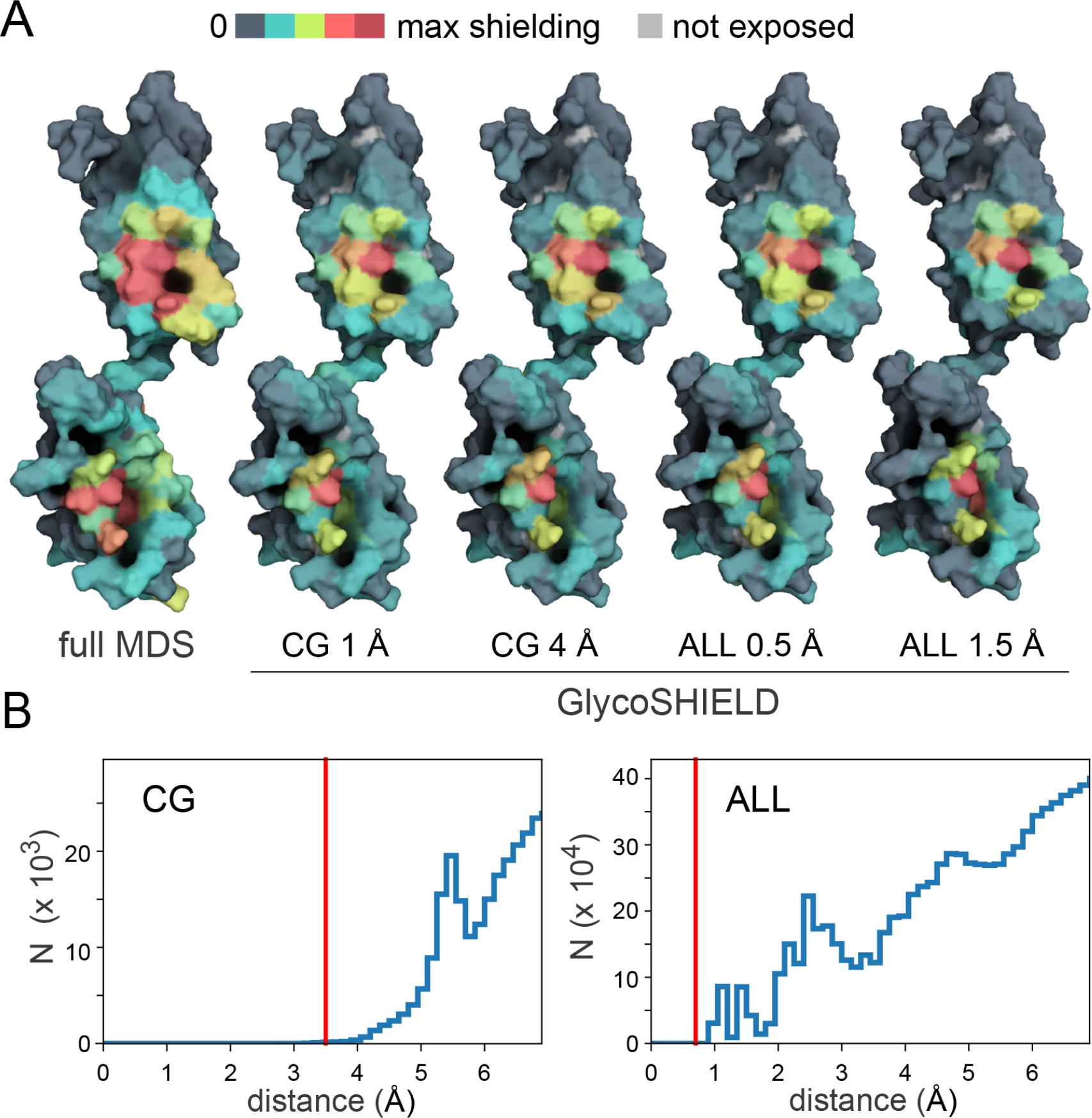
EC4-EC5 shielding estimated by MDS and GlycoSHIELD. **A)** 3D-heatmaps of protein shielding (i.e. glycan-dependent reduction of solvent accessible surface area, see Methods) by glycan conformers computed by MDS or GlycoSHIELD for multiple exclusion distances between protein α-carbons and glycan ring oxygens (coarse grain or CG) or all protein and glycan atoms (ALL). **B)** Left panel: distribution of glycan oxygen atoms at increasing distance from protein α-carbons (CG) calculated from 1 μs trajectory at 1 ns intervals; N: occurrence counts; right panel: distribution of glycan atoms at increasing distance from protein atoms. Threshold values selected for shield reconstructions (e.g., Figure 1C) are indicated in red. In A, note the good agreement of shielding heatmaps derived from MDS and GlycoSHIELD.

**Supplemental Figure S3.**
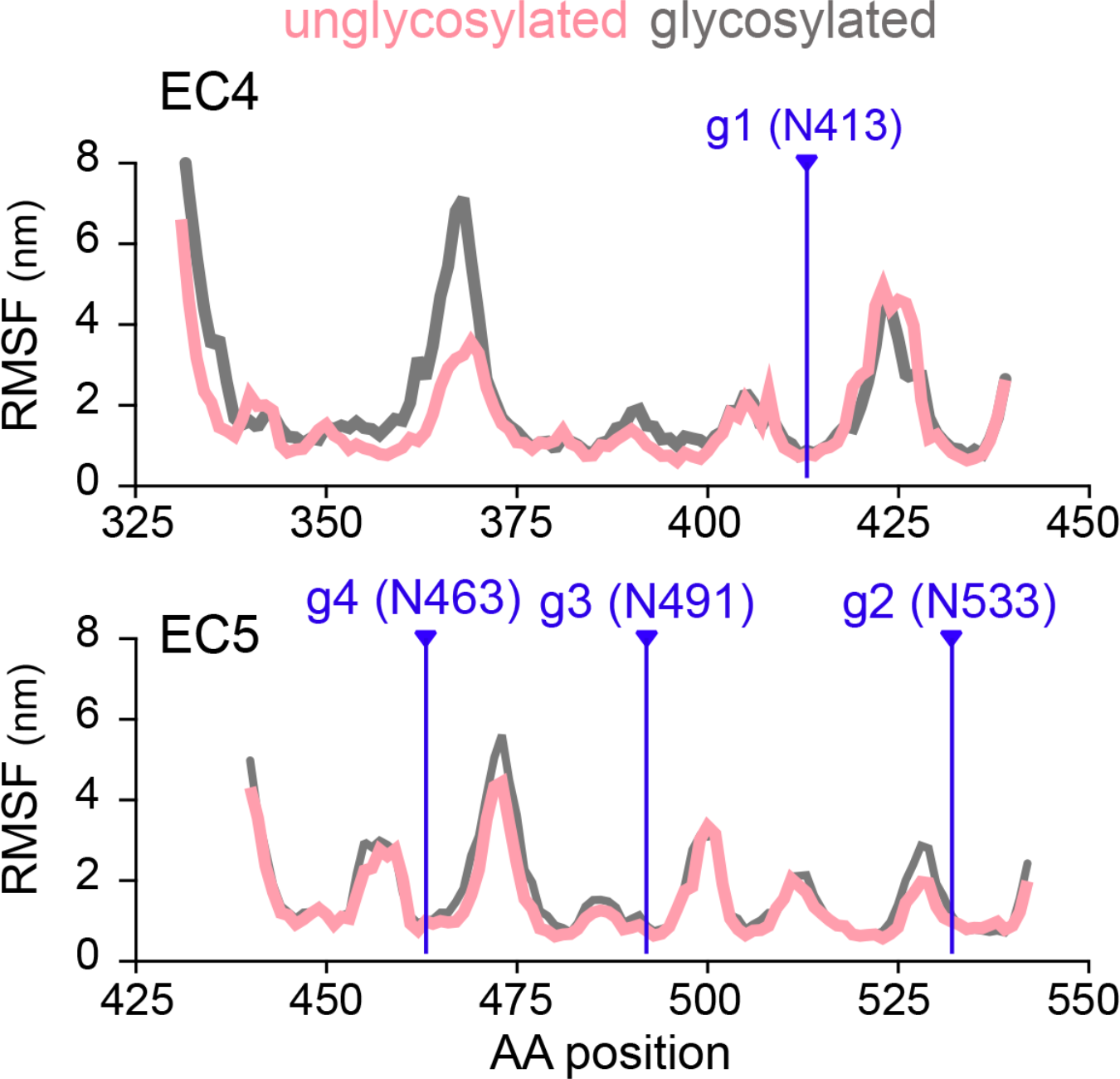
N-glycans have a low impact on protein motion within stable EC domains. Root mean square fluctuations (RMSF) of C alpha atoms over time for EC4 or EC5 domains in unglycosylated and glycosylated proteins (glycosylation sites are indicated by vertical lines). In A and B, note the low impact of glycans on protein motion within each domain.

**Supplemental Movie S2. Conformational change of unglycosylated and glycosylated EC4-EC5 over the entire simulation.** Glycans are not represented. Unglycosylated and glycosylated EC4-EC5 are shown on the left and the right panel, respectively.

**Supplementary Figure S4.**
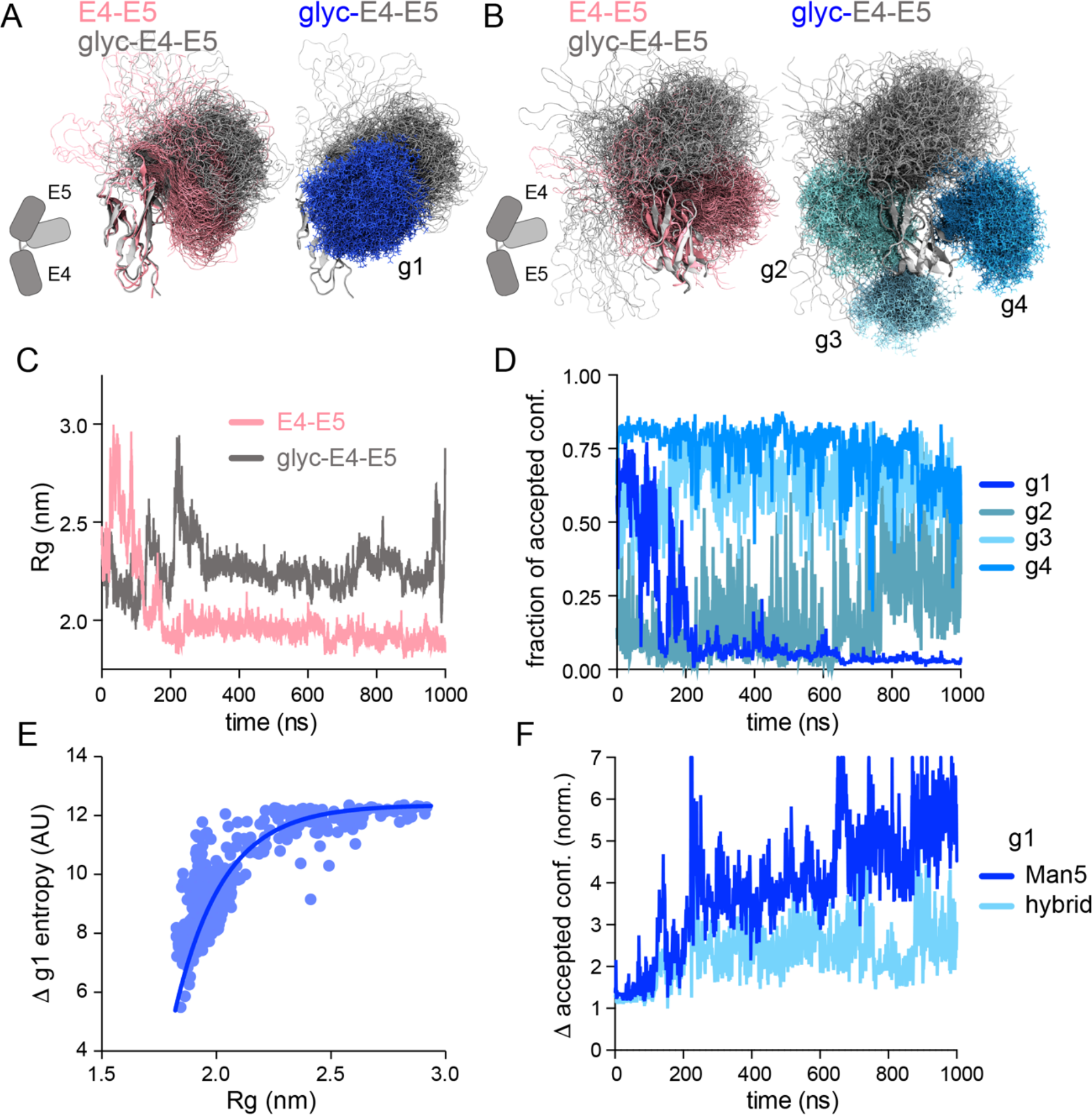
GlycoSHIELD provides realistic predictions on EC4-EC5 structural dynamics. **A-B)** Movements of N-cadherin EC4 and EC5 domains in relation to one-another in unglycosylated EC4-EC5 or glycosylated EC4-EC5 (glyc-EC4-EC5). Shown are positions of protein backbones (pink or grey) and glycans (blue) displayed as ribbons (fixed EC4 or EC5 domains) or superposed lines (moving glycans or protein backbone). Glycans overlapping (g1) or non-overlapping with possible protein positions (g2-g4) are shown in dark and lighter blue, respectively. **C)** Variation of the gyration radius of EC4-EC5 (pink) or glyc-EC4-EC5 (grey) over the simulation. Note the adoption of a compact (reduced Rg values) and a more elongated conformation by EC4-EC5 and glyc-EC4-EC5, respectively. See also Supplementary Fig.S3 and Movies S1 and S2. **D)** Fraction of accepted GlycoSHIELD conformers for g1-g4 (same colors as in B-E) grafted on the conformers from the EC4-EC5 simulation. Note the strong correlation between the compaction of the protein (reduced Rg values in the pink plot in C) and clashing with glycans (reduced fraction values for glycan shown in dark blue). **E)** Corresponding variation of estimated partial entropy of g1 (κ_B_ ln[Ω], with κ_B_ Boltzmann constant and Ω conformer number) as a function of protein radius of gyration, showing increasing glycan steric constraints for Rg values below 2.25 nm. **F)** Normalized variations of accepted conformers for g1 with Man5 (high-mannose) or Man5_Gal1_Fuc1 (hybrid) sugars as compared to original Fuc1_Neu4 (complex) structure, showing the effect of glycan type on glycan steric constrains.

**Supplemental Figure S5.**
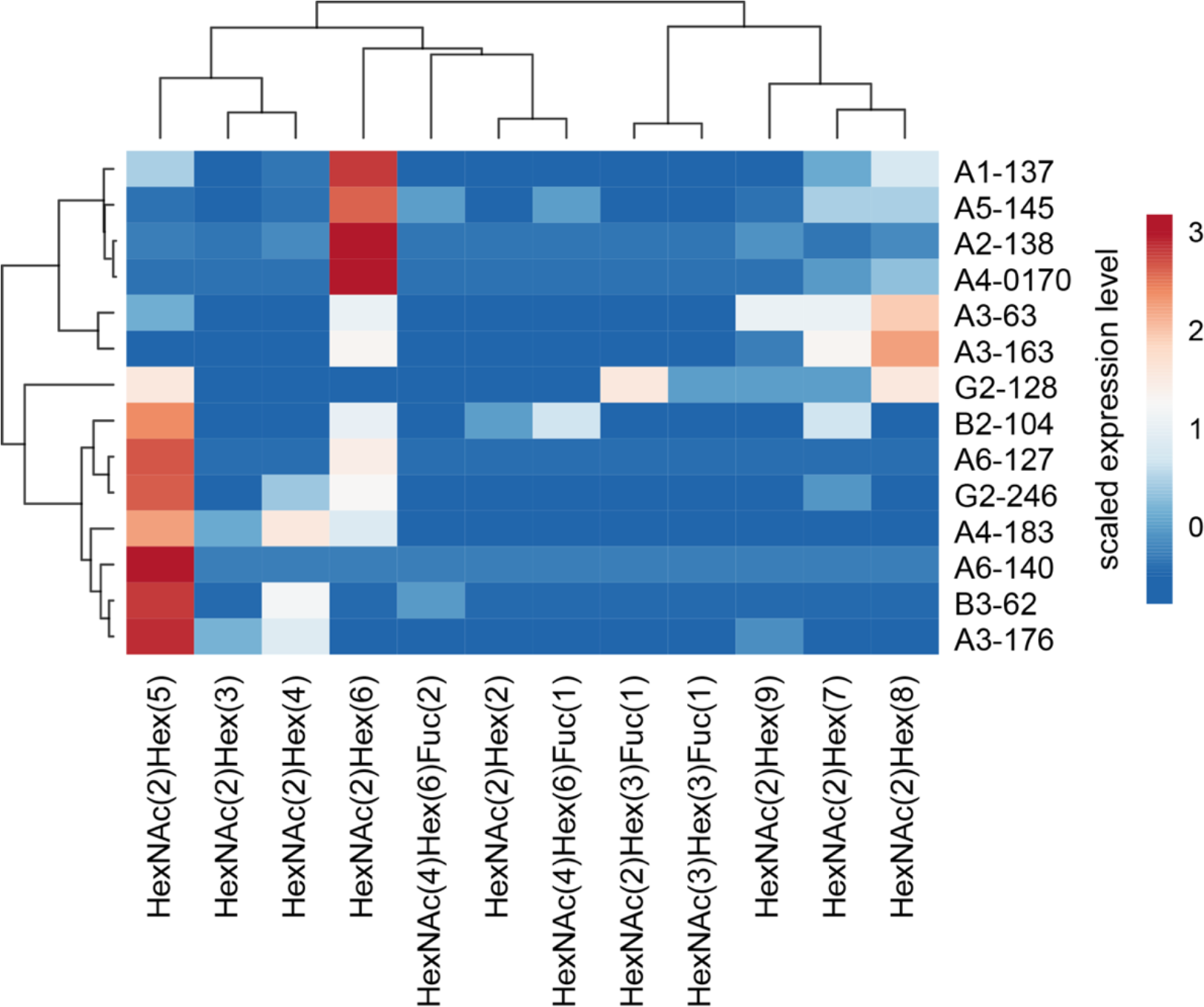
GlcNac2-Man5 (Man5) is the most abundant N-glycan of GABAA receptors in the mouse brain. Clustering heatmap (correlation matrix) showing the variability and abundance of N-glycans detected by mass spectrometry on GABA_A_ receptor subunits in the mouse brain. Generated from raw data published in Riley, N.M. et al., *Nat Commun* **10**, 1311 (2019). doi.org/10.1038/s41467-019-09222-w. Labels on the right side indicate subunit type and glycosylation sites (e.g. B3-63 is β3-subunit N63). Note the marked over-representation of Man5 [HexNac(2)-Hex(5)].

**Supplemental Figure S6.**
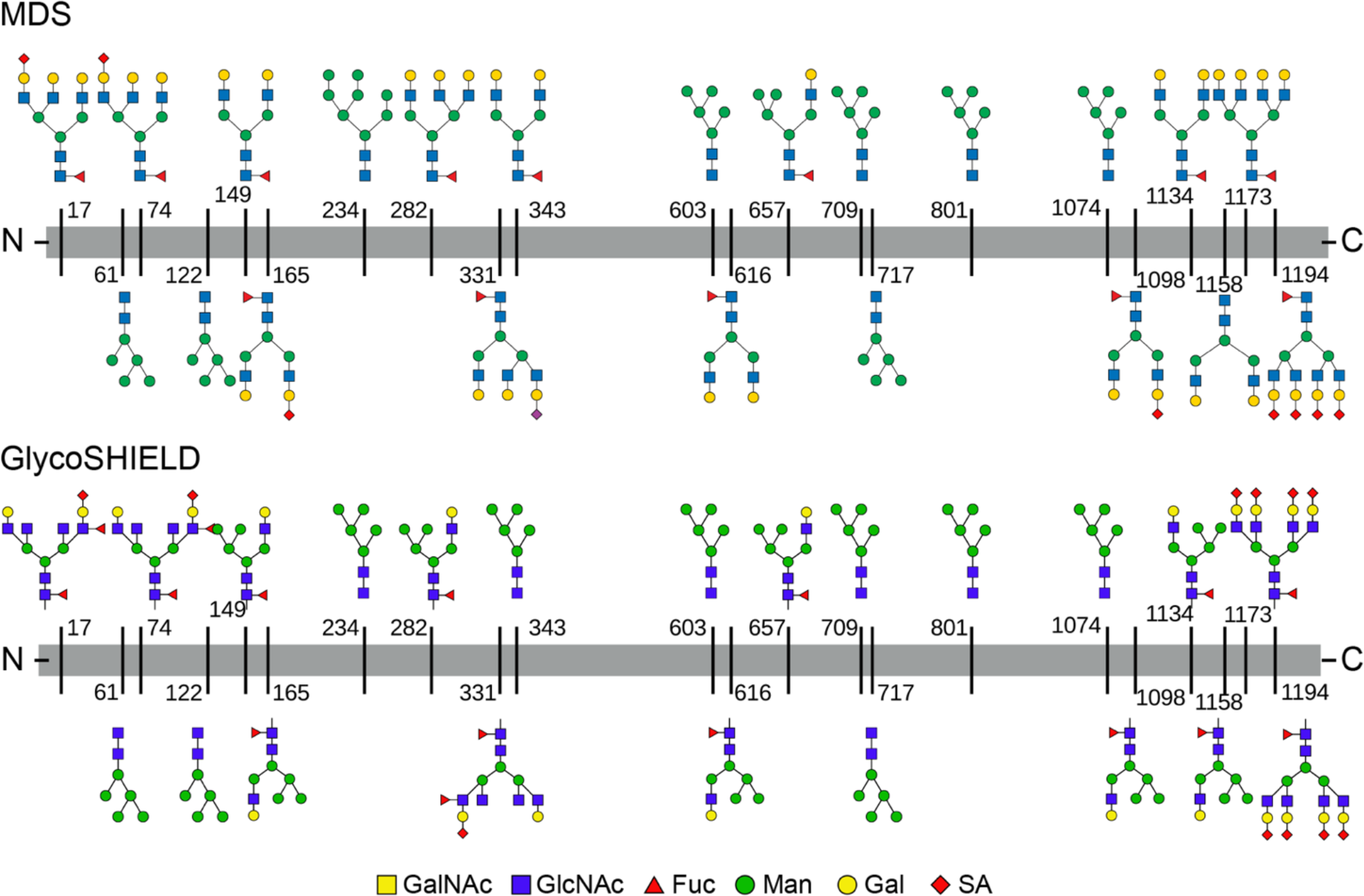
Glycosylation profile of S-protein used for MDS and GlycoSHIELD (see Figures 3A)

**Supplemental Figures S7.**
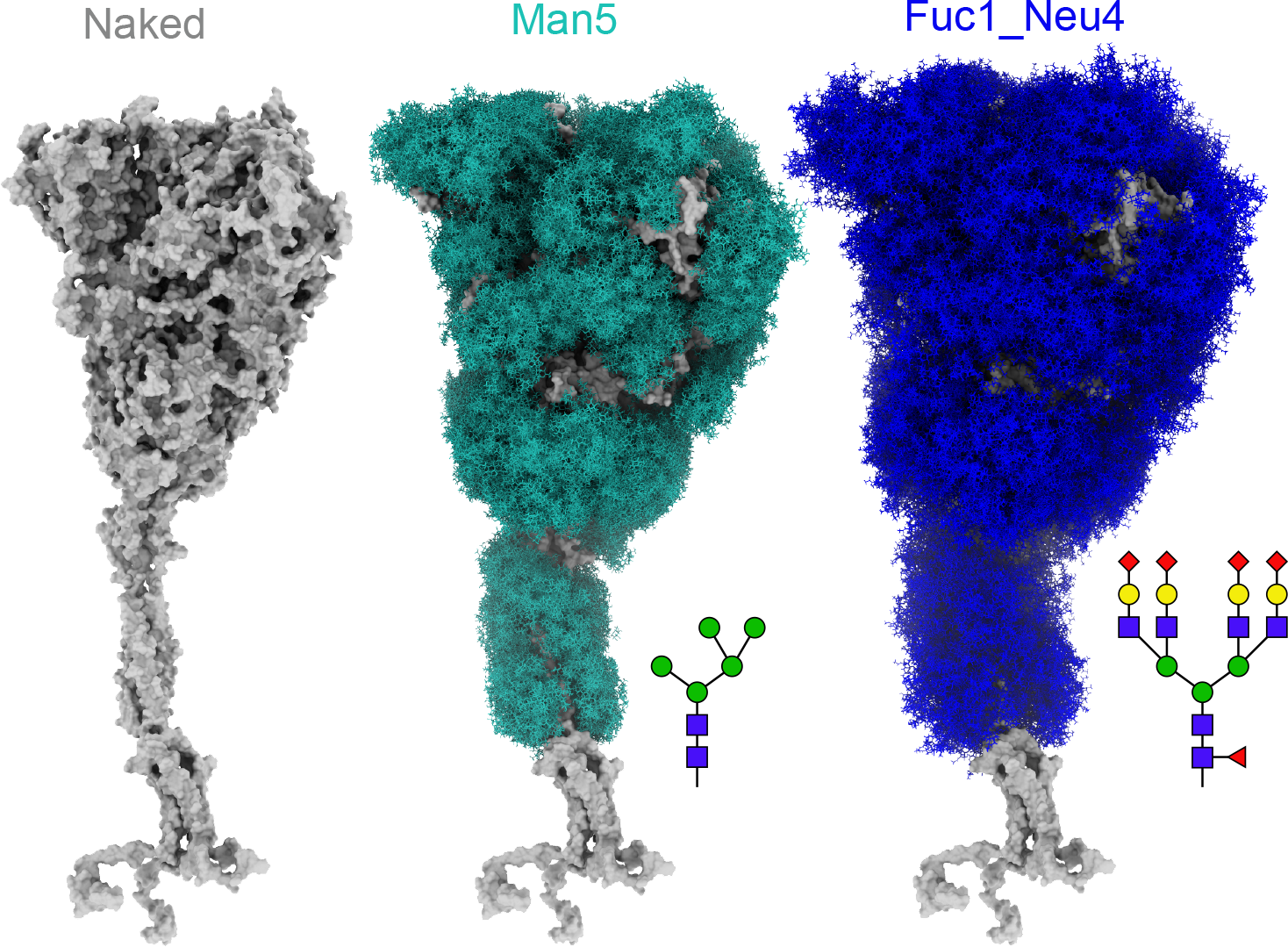
S-protein modelled without glycans or with GlycoSHIELD with all Man5 or all Fuc1_Neu4 N-glycans. Note the increased span of the Fuc1_Neu4 shield compared to Man5.

**Supplemental Figures S8.**
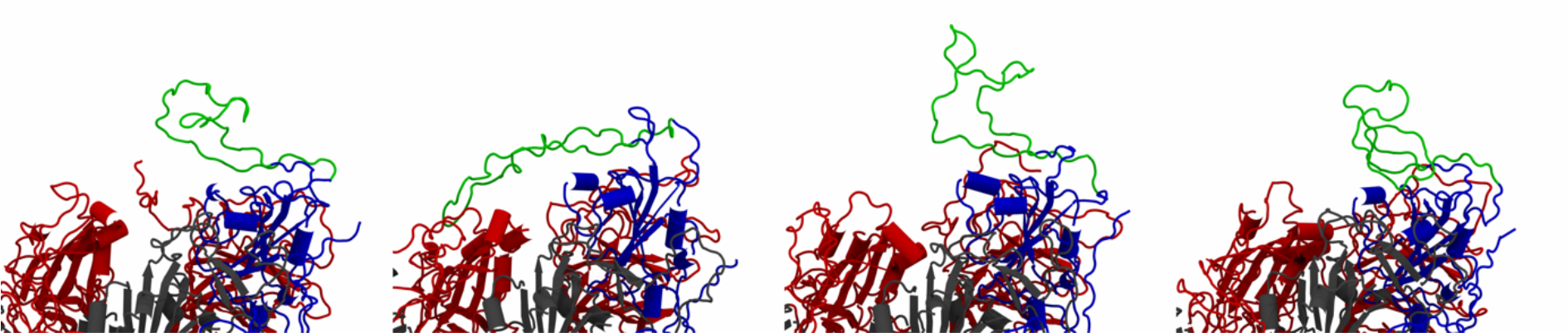
Mobility of the loops on the RBD. Representative snapshots of S-protein (blue, red and gray, cartoon representation) were taken from the 10 μs simulation and flexible loop of the RBD (residues 445-495) was highlighted in green.

## Notes

https://github.com/GlycoSHIELD-MD/GlycoSHIELD-0.1

https://zenodo.org/record/5083355

### Competing Interest Statement

The authors have declared no competing interest.

